# Direct RNA nanopore sequencing reveals rapid RNA modification changes following glucose stimulation of human pancreatic beta-cell lines

**DOI:** 10.1101/2025.06.12.659352

**Authors:** Logan Mulroney, Henry J. Taylor, Angela Lee, Amy J. Swift, Mihail Zdravkov, Lori L. Bonnycastle, Shelise Y. Brooks, Brian N. Lee, Tomas Fitzgerald, Narisu Narisu, Leslie G. Biesecker, Michael R. Erdos, Francesco Nicassio, Ewan Birney, Francis S. Collins, D. Leland Taylor

## Abstract

RNA modifications are critical regulators of gene expression and cellular processes; however, the epitranscriptome is less well studied than the epigenome. Here, we studied transcriptome-wide changes in RNA modifications and expression levels in two human pancreatic beta-cell lines, EndoC-BH1 and EndoC-BH3, after one hour of glucose stimulation. Using direct RNA nanopore sequencing (dRNA-seq), we measured N6-methyladenosine (m6A), 5-methylcytosine (m5C), inosine, and pseudouridine concurrently across the transcriptome. We developed a differential RNA modification method and identified 1,697 differentially modified sites (DMSs) across all modifications. These DMSs were largely independent of changes in gene expression levels and enriched in transcripts for type 2 diabetes (T2D) genes. Our study demonstrates how dRNA-seq can be used to detect and quantify RNA modification changes in response to cellular stimuli at the single-nucleotide level and provides new insights into RNA-mediated mechanisms that may contribute to normal beta-cell response and potential dysfunction in T2D.

## Introduction

Chemical modifications of RNA molecules are important regulators of gene expression, whereby “writer” and “eraser” enzymes alter the molecular structure of nucleotides in RNA and lead to post-transcriptional regulatory functions^1^. To date, studies have described more than 160 distinct naturally occurring RNA modifications^2^, of which 13 have been documented in mammalian mRNA^3^—each with specific downstream mechanisms and disease-relevant implications spanning cancer, birth defects, and neurological disorders^1,4^.

Historically, research in RNA modifications has faced several limitations. Many of the methods used to measure RNA modifications require laborious, RNA-modification-specific biochemical assays, such as RNA modification immunoprecipitation or chemical reactions that have unique interactions with target RNA modifications^5^. Furthermore, it is challenging to combine assays to detect multiple RNA modifications simultaneously due to conflicting biochemical reactions or assay detection methods. Thus, separate samples are often required for each modification-specific assay. The labor and other experimental constraints have made it challenging to study multiple RNA modifications simultaneously within model cell lines or primary patient samples.

Direct RNA nanopore sequencing (dRNA-seq) by Oxford Nanopore Technologies (ONT) is a single-molecule, long-read sequencing method that reads native RNA molecules without creating complementary DNA (cDNA)^6^. Without the cDNA conversion step, RNA modifications are preserved during sequencing and can be identified using computational models trained on their unique signatures from the nanopore sequencer^7,8^. In theory, dRNA-seq technology could potentially detect all possible RNA modifications in a nucleotide-specific manner and greatly expand the ability to characterize the role of post-transcriptional gene regulation in disease^9^. To date, ONT has released models to detect four modifications—m6A, m5C, inosine, and pseudouridine—during basecalling, and although there have been studies in resting cell lines or baseline samples, the application of such technologies in disease-relevant contexts remains limited.

Type 2 diabetes (T2D) is a leading cause of global morbidity and mortality^10^. Although the role of pancreatic islet beta-cell dysfunction is a widely recognized feature of T2D pathophysiology, the underlying molecular mechanisms leading to this dysfunction remain unclear^11^. Recent studies exploring m6A levels in pancreatic islets suggest an important role of RNA modifications in beta-cell response to glucose and in T2D^12,13^. De Jesus et al. identified decreased m6A levels on transcripts for insulin-regulating genes in T2D islets and showed that reduced m6A “writer” levels led to impaired insulin secretion in a human beta-cell line and early onset diabetes in mice^12^. Similarly, Bornaque et al. observed a sharp decrease of m6A methylation in MIN6 mouse beta-cells and primary human islets upon high glucose stimulation^13^. Despite this evidence of m6A-mediated islet response and dysfunction in T2D, there have been limited efforts to characterize the remaining modifications across the beta-cell transcriptome in T2D-relevant conditions.

Here, we exposed two human beta-cell lines, EndoC-BH1^14^ and EndoC-BH3^15^, to euglycemic and hyperglycemic conditions for one hour, performed dRNA-seq, quantified gene expression, transcript expression, and four different RNA modification levels, and tested all these aspects of gene expression for associations with glucose exposure. We identified 1,697 differentially modified sites (DMSs), 209 differentially expressed genes, and 121 differentially expressed transcripts. We found that the DMSs for m6A, m5C, and inosine were enriched in T2D genes, but there was no T2D enrichment for the differentially expressed genes and transcripts. Additionally, the DMSs were largely independent from the expression changes. These results provide additional molecular insights into beta-cell response and demonstrate how dRNA-seq can be used to interrogate multiple RNA modifications simultaneously, allowing for a more complete understanding of molecular interactions and disease-relevant mechanistic insights.

## Results

### Native RNA sequencing in human beta-cell lines after glucose stimulation

We cultured EndoC-BH1 (N = 3) and EndoC-BH3 (N = 4) cell lines in euglycemic (2.8 mM) and hyperglycemic (15 mM) conditions for one hour (Fig. 1A). We measured secreted insulin (Methods) and found that insulin secretion was induced, as expected in normal beta-cell function (*P*-value<0.05, Fig. 1B). We harvested poly(A) RNA and performed dRNA-sequencing using RNA004 PromethION flowcells from ONT (Methods). Across the native RNA samples, we obtained an average of 18.5 million reads per flowcell with an average N50 of 1.7 kb (Supplementary Table 1). In addition, for several samples, we created a modification-free reference transcriptome using *in vitro* transcription (IVT) from cDNA (Fig. 1A; Methods)^16^. We sequenced the IVT-derived RNA using the same approach as the native RNA and obtained an average of 5.3 million reads per flowcell from the IVT RNA samples (Supplementary Table 1).

**Fig. 1.**
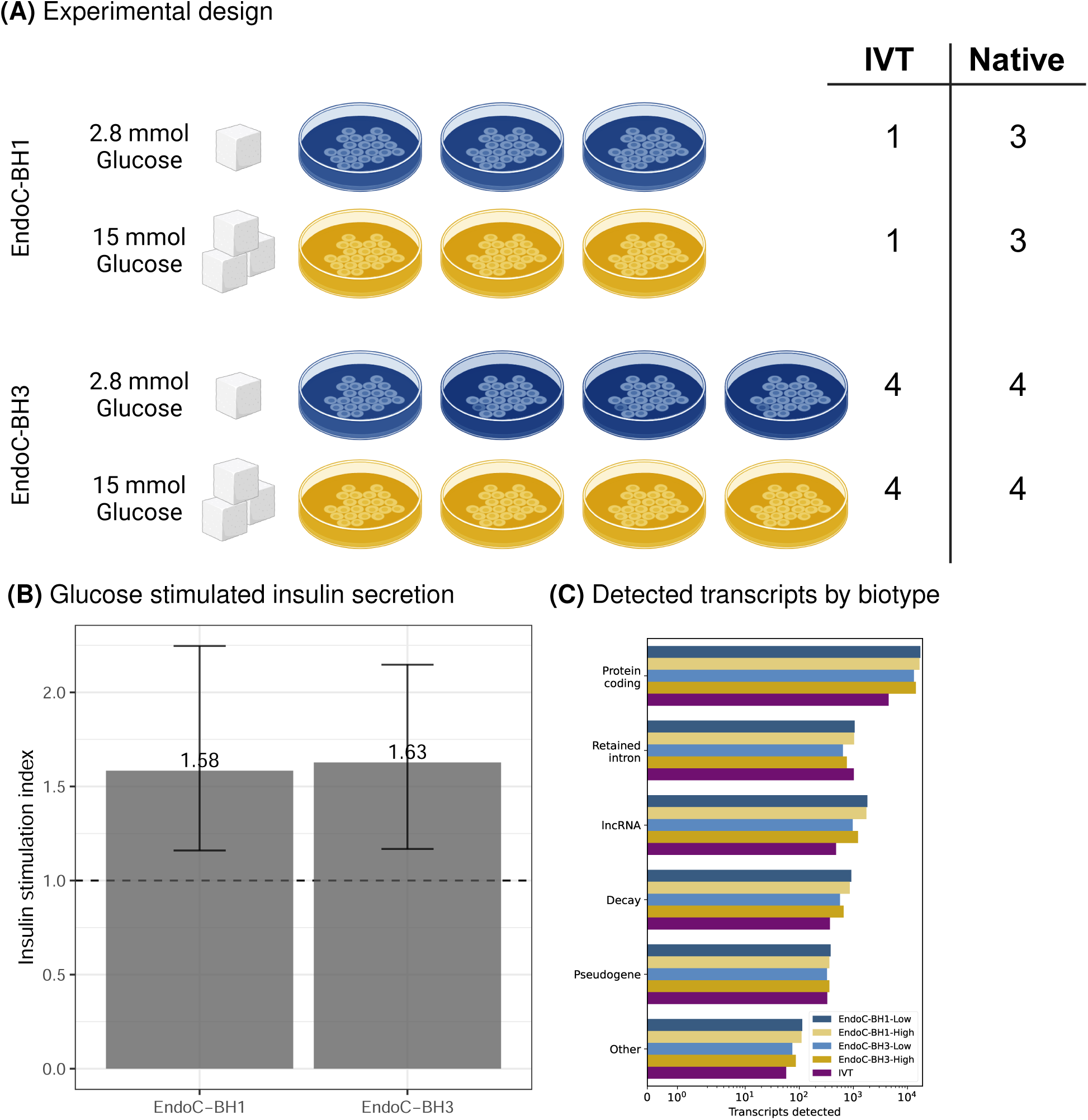
Study summary. (A) EndoC-BH1 (top) and EndoC-BH3 (bottom) were cultured in low glucose condition (2.8 mM, blue) or high glucose condition (15.0 mM, yellow). RNA was extracted from each culture and used to prepare native dRNA-seq libraries or *in vitro* transcribed (IVT) dRNA-seq libraries. (B) Insulin stimulation index (y-axis) across cell lines (x-axis) after one hour of glucose exposure. (C) Detected transcripts (x-axis) by biotype (y-axis).

One advantage of long-read RNA sequencing over short-read sequencing is the ability to capture full-length transcripts. Using the GRCh38 reference genome and gencode v46 reference transcriptome, we aligned the reads and quantified isoform-resolved transcript levels (Methods)^17^. To control for potential technical artifacts, we filtered for transcripts with at least three mean reads across replicates and identified 36,200 unique isoforms generated from 16,469 unique genes. As expected due to the poly(A) selection, 24,383 (67.4%) of the 36,200 isoforms were protein coding genes (Fig. 1C).

We used the ONT modification tool, remora, during basecalling with dorado to detect RNA modifications^9,18^ in the native and IVT samples (Methods). At the time of analysis, remora was capable of calling m6A, m5C, inosine, or pseudouridine modifications due to limited training data to develop other modification-specific models^9^. We identified 47,555,270 transcriptome positions that spanned 36,319 unique transcript isoforms with at least one modified nucleotide and with at least 20 average reads across the native sample replicates. Although we found that the modification proportion for each modified site was strongly correlated between the two cell lines (minimum r^2^>0.80 across all modifications; Supplementary Fig. 1), we also found that there were several likely false positive modifications detected in the IVT samples (Supplementary Fig. 2). We filtered likely false positive modifications from the native RNA by including modification sites with at least 5% modification proportion in the native samples, less than 5% modification proportion in the IVT samples, and at least 20 mean reads across native replicates (Methods). Using these filtering conditions, we identified 2,706,493 total modified sites: 1,468,238 m6A, 495,376 m5C, 195,097 inosine, and 547,782 pseudouridine sites.

### Differentially modified sites for known RNA modifications

We tested for RNA modifications associated with glucose exposure within each cell line using binomial regression and meta-analyzed the results (Methods). We identified 1,337, 321, 21, and 18 DMSs for m6A, m5C, inosine, and pseudouridine, respectively (false discovery rate [FDR]<5%; Fig. 2A). Of the 28,565 genes we measured, we found glucose-associated modifications on 1,050 unique genes and 1,133 unique transcripts, highlighting a widespread modification response to glucose. We overlapped these genes with well-established T2D genes from the T2D knowledge portal^19,20^ and observed an enrichment of T2D-related genes across m6A, m5C, and inosine, but not pseudouridine (FDR<5%, hypergeometric test; Fig. 2B; Supplementary Fig. 3).

**Fig. 2.**
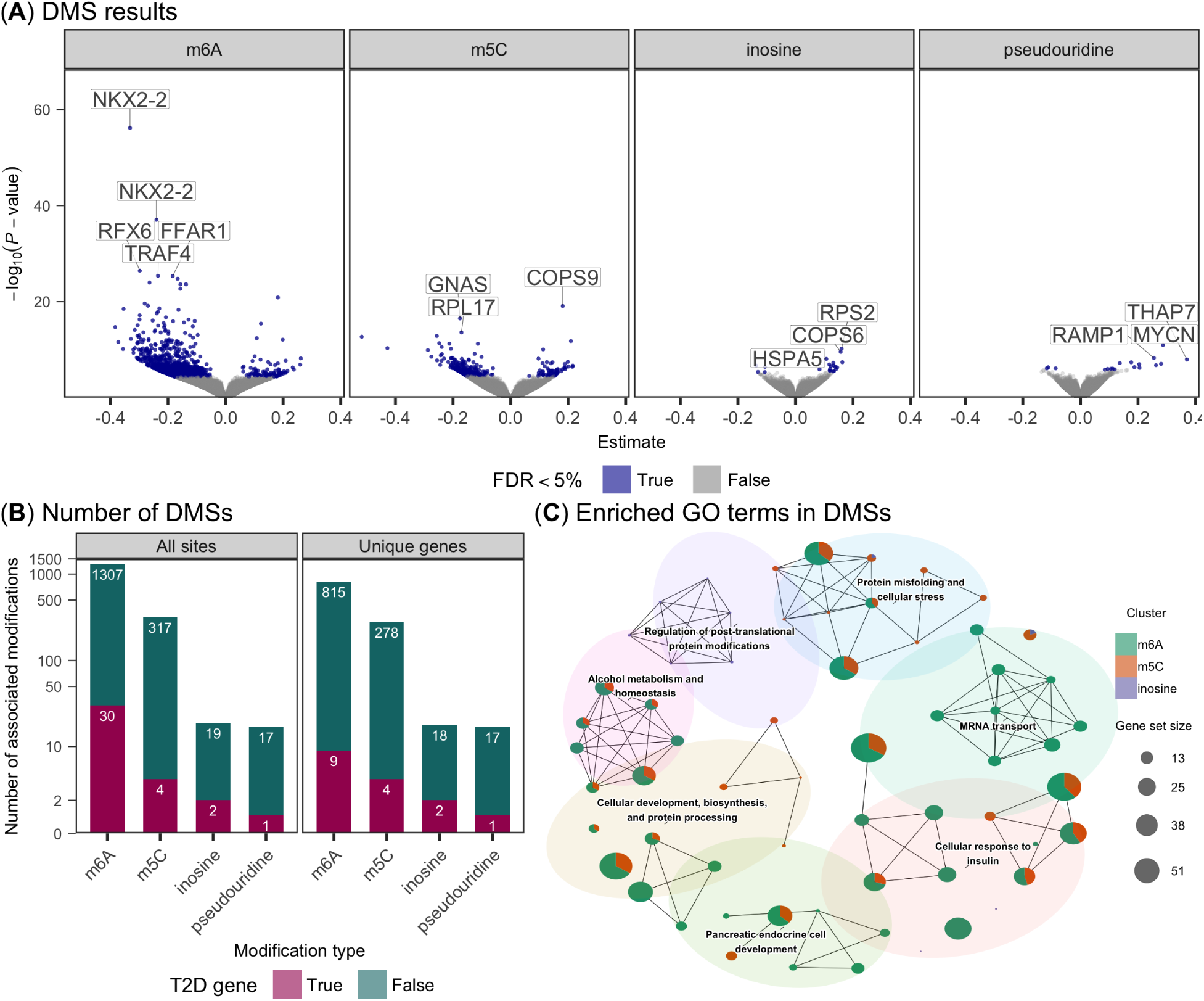
Differential RNA modification results. (A)The estimated effect size (x-axis) and *−log*_10_(*P*-value) (y-axis) for modification sites (points) after high glucose exposure across RNA modifications (facets). A positive effect size corresponds to an increased proportion of modified counts in the high glucose state. Blue points depict FDR*<* 5% and a subset of genes that contain the DMSs are labeled. (B) Bar plot of the number DMSs (left) or genes with at least one DMS (right), colored by T2D gene status. (C) Network of GO terms enriched (FDR*<* 5%) in differentially modified sites. Each node is a different GO term with the proportion of each DMS represented by the pie chart. The node size is scaled by the number of genes overlapping each GO term. Edges connect semantically similar nodes, which are also clustered based on similarity.

To explore the biological processes underlying the modified genes, we performed an enrichment analysis of biological process gene ontology (GO) terms using genes with at least one DMS and clustered enriched GO terms based on semantic similarity (Methods). We identified 62 enriched gene sets that clustered in seven putative functional groups (FDR<5%; Fig. 2C). We found several terms related to cellular response to insulin, pancreatic cell development, and metabolism—underscoring the importance of RNA modifications in response to glucose in EndoC-BH1 and EndoC-BH3 lines (Fig. 2C).

We identified several T2D-related genes with multiple modifications, two of which we highlight. *PDX1* is an essential transcription factor for beta-cell development^21^. In the primary *PDX1* transcript, *PDX1-201*, we found one pseudouridine and two m6A DMSs (Fig. 3A). Notably, the two m6A DMSs identified in the 3’ UTR of PDX1-201 are consistent with regions of differential m6A peaks from prior observations in islets from patients with normal glucose tolerance and T2D^12^. We also identified multiple DMSs in *NKX2-2*, another transcription factor that is essential for beta-cell cell development and maintenance in humans^22,23^. We detected RNA modifications at several positions along the *NKX2-2-201* transcript body, including one inosine and nine m6A DMSs (Fig. 3B). Of the nine m6A modifications, two were near the stop codon, as was expected based on previous studies of m6A localization^24^. Surprisingly, the m6A DMS with the strongest glucose response effect was located near the start codon.

**Fig. 3.**
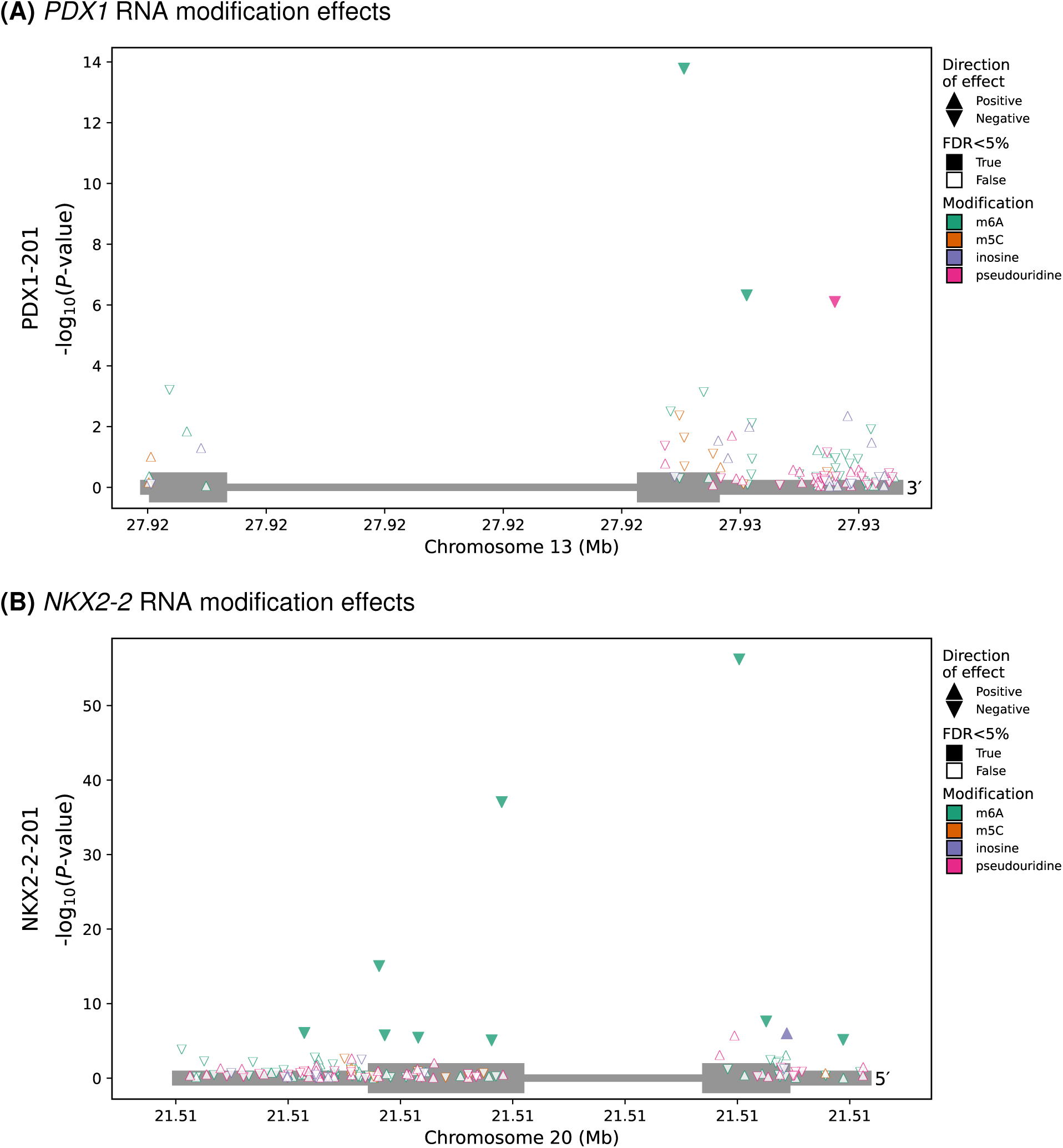
RNA modification results at transcript resolution. RNA modification sites considered (points) along with their genomic coordinates (x-axis) in relationship to the detected transcripts of *PDX1-201* (A) and *NKX2-2-201* (B) (grey boxes) and the differential modification *−log*_10_(*P*-value) (y-axis). Direction of effect indicated by triangle orientation, where an upward triangle indicates a positive effect (i.e., increased proportion of modified counts in the high glucose condition). Colors depict the RNA modification. Fill status indicates FDR*<* 5%.

### Differentially expressed genes and transcripts

We characterized the transcriptional profile of the cell lines after the one-hour glucose exposure, performing differential gene expression (DGE) and differential transcript expression (DTE) analysis. Similar to the RNA modification analysis, we tested genes and transcripts within the two cell lines and meta-analyzed common features (Methods). We found 209 genes and 121 transcripts that were differentially expressed (FDR<5%; Fig. 4A). In contrast to the DMS results, we did not identify an enrichment of T2D genes (*P*-value≥0.05, hypergeometric test; Methods).

**Fig. 4.**
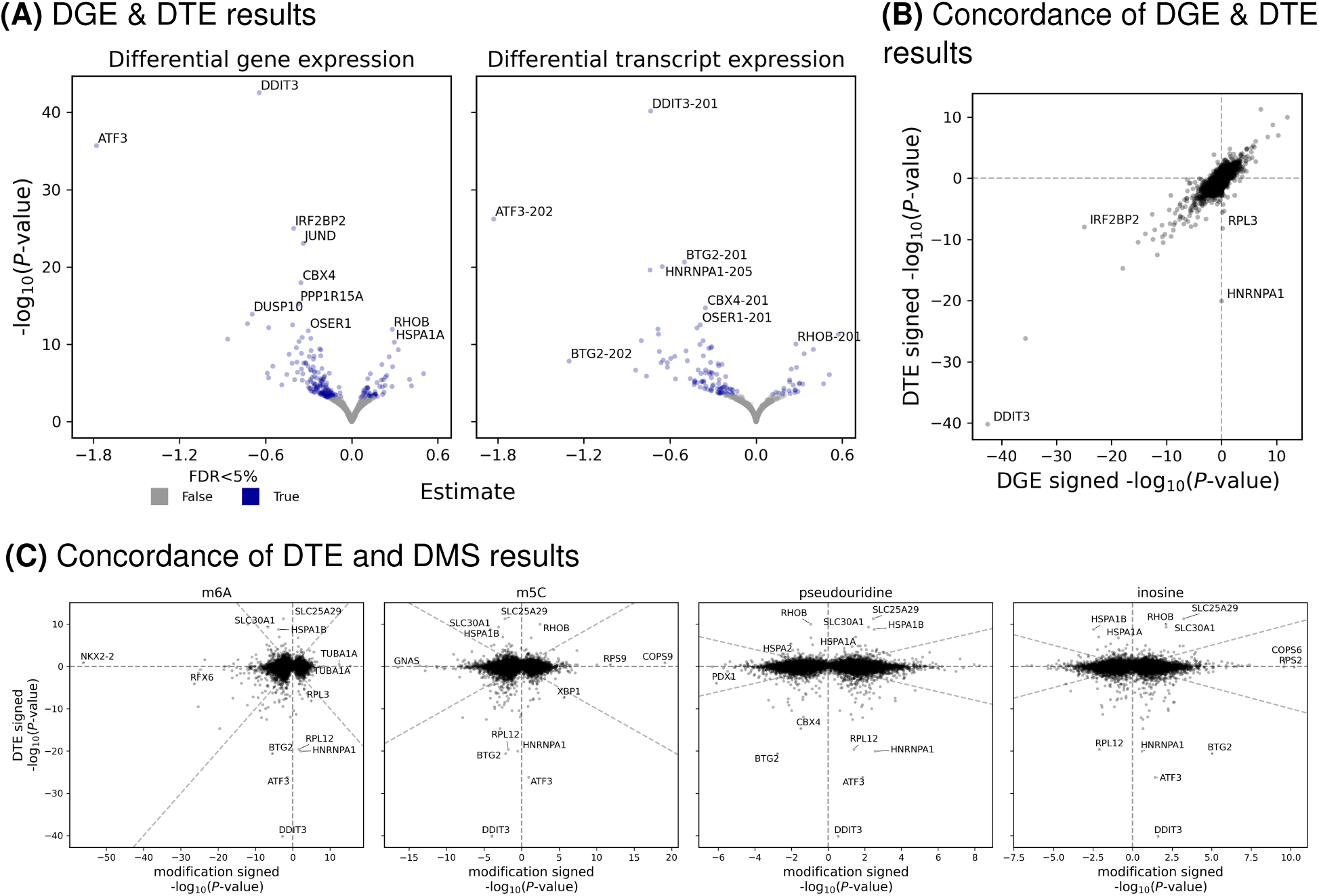
Differential gene and transcript results. (A) The estimated effect size (x-axis) and *−log*_10_(*P*-value) (y-axis) for gene or transcript expression (points, facets) after high glucose exposure. Blue points depict FDR*<* 5%. (B) Comparison of the signed *−log*_10_(*P*-value) for genes (x-axis) and the transcript for the corresponding gene with the smallest *P*-value (y-axis). (C) Comparison of the signed *−log*_10_(*P*-value) for transcripts (y-axis) and the signed *−log*_10_(*P*-value) of the RNA modification (facets) for the corresponding transcript with the smallest *P*-value (x-axis).

We considered the concordance between DTE and DGE signals and found strong consistency between the DGE signals and the DTE signals with the smallest *P*-value for the respective gene (Fig. 4B), which comports with observations that gene expression is often dominated by a single isoform (reviewed in Tress et al.^25^). There were two notable exceptions, *HNRNPA1* and *RPL3*, where we identified a DTE signal but no detectable DGE change. For both of these associations, a single transcript showed changes in response to glucose stimulation, but that signal was masked by other invariant transcripts when considering total gene expression (Supplementary Fig. 4).

We tested for biological processes GO terms enriched in both our DGE and DTE results and clustered the gene sets based on semantic similarity (Methods). We identified 54 enriched gene sets that clustered in seven putative functional groups that were associated with glucose exposure (FDR<5%). As with the DMS results, many of these terms were associated with cellular metabolism and glucose response, suggesting an initial transcriptional response even after one hour of stimulation (Supplementary Fig. 6).

Finally, we considered the concordance between the DTE and DMS signals. We observed very little signal concordance between the DTE and DMS associations across the modification types (Fig. 4C)—only 41 differentially expressed transcripts contained at least one DMS (33.9% of DTEs). Combined, these results suggest that differences in transcript expression were not drivers of the observed differences in RNA modifications. Indeed, of the known writers and erasers for the considered RNA modifications, none were differentially expressed (Supplemental Table 2).

## Discussion

RNA modifications are highly dynamic chemical alterations to ribonucleotides that play an important role in the cellular response to stimuli^1^. Thirteen of the greater than 160 RNA modifications described have been identified in mammalian mRNA^2,3^. However, simultaneous profiling of multiple RNA modifications transcriptome-wide in disease-relevant contexts has been limited by technological constraints. Recent advances in dRNA-seq and related computational tools have made it possible to quantify isoform-resolved, single-nucleotide modifications at a transcriptome-wide scale across many samples^6,9,18^. Here, we use dRNA-seq to perform a transcriptome-wide analysis of differentially modified sites, differentially expressed genes, and differentially expressed transcripts in two human beta-cell lines after a one hour glucose challenge and evaluate our findings in the context of T2D.

We called RNA modifications using ONT’s remora to identify four specific modifications: m6A, m5C, inosine, and pseudouridine. To control for false positive modification calls, we created a modification-free IVT dataset. The IVT reference served as a critical quality control step and allowed us to identify and remove potentially false positive modification sites called by remora: 13.7% of m6A sites, 6.9% of m5C sites, 5% of pseudouridine sites, and 1.3% of inosine sites (Supplementary Fig. 2). This step will likely become less critical as RNA modification detection models improve over time, but for now measures to curtail false positives are required.

Focusing on the modifications, we found that m6A represented 78.8% (1,337) of the DMSs, consistent with previous literature that describes an m6A-mediated effect on glucose response in beta-cells (Fig. 2A)^12,13^. Of the m6A-modified genes, we found an enrichment for T2D-related genes, such as *PDX1* and *NKX2-2*, consistent with prior observations^12^. Although most m6A sites are found in the 3′ UTR near the stop codon^24^, we found a number of differentially modified m6A sites near the start codon (Fig. 3B). It has been documented previously that m6A can alter RNA splicing, decay, and translation through interactions with YTH-domain m6A reader proteins^26^. Given the lack of concordance between the differential expression changes and the differential m6A sites that we found, we speculate that the m6A changes act through a pathway different than RNA decay (Fig. 4C). Beyond m6A, we also identified 321, 21, and 18 glucose associations for m5C, inosine, and pseudouridine, respectively. To our knowledge, this study is the first description of m5C, inosine, and pseudouridine in beta-cell glucose response. Akin to m6A, the DMSs signal for these other RNA modifications is primarily independent from the observed transcription expression differences (Fig. 4C). Future studies within beta-cells will be required to understand whether these changes causally mediate normal beta-cell response to glucose stimulation. In addition, analysis of additional time points will also help determine the molecular consequences of the RNA modification changes on expression or translation changes in response to glucose stimulation.

In summary, our work represents one of the first large-scale, transcriptome-wide efforts to simultaneously characterize multiple different RNA modifications in cellular response to a physiologically relevant stimulus. We identify rapid and disease-relevant alterations in several RNA modifications following glucose stimulation of beta-cells. We anticipate that RNA modifications may also be important for a variety of other diseases in various cell types. Our study serves as a model for future studies of differential RNA modifications.

## Materials and Methods

### Cell culture, glucose stimulation, and measurement of secreted insulin

We cultured EndoC-BH1^14^ and -BH3^15^ cells using flasks, plates, or dishes coated with 500X DMEM/F-12 + GlutaMAX™ (Thermo Fisher Scientific, TFS) containing 1X Fibronectin, 5X Penicillin-Streptomycin, and 5X Extracellular Matrix (Sigma), and incubated them for at least 1 hour at 37°C and 5% CO₂. To prepare the culture medium, we supplemented Advanced DMEM/F-12 (TFS) with 50 μM 2-mercaptoethanol, 2% BSA (Equitech-Bio), 6.7 ng/mL sodium selenite (Sigma), 5.5 μg/mL transferrin (Sigma), 2 mM GlutaMAX (TFS) and 10 mM nicotinamide (VWR), and sterilized it using a 0.22 μm filter (Corning). For EndoC-BH3 cells, we added puromycin (TFS) to a final concentration of 10 μg/mL. We prepared trypsin neutralization media by mixing 80% PBS (TFS) with 20% heat-inactivated FBS (Bio-Techne) and filtered it using a 0.22 μm filter (Corning).

For passaging, we prewarmed 0.05% Trypsin (TFS) and treated cells for 2 minutes at 37°C. We neutralized trypsin with an equal volume of neutralization media, pipetted to dissociate cells, and added culture media at 3x volume. We counted cells using a NucleoCounter® NC-3000™ (Chemometec), seeded them at 80,000-90,000 cells/cm², and passaged them weekly. For EndoC-BH3 cells, we maintained puromycin selection until we induced CRE-mediated excision of immortalizing transgenes by adding 1 μM tamoxifen (TAM, Sigma) for three consecutive weeks, followed immediately by glucose-stimulated insulin secretion (GSIS). Cell expansion and TAM treatment required approximately 8 weeks. We cultured EndoC-BH1 cells using the same conditions, excluding TAM and puromycin, and expanded them over 4 weeks.

To perform GSIS, we seeded EndoC-BH1 cells and TAM-treated EndoC-BH3 cells in 6-well plates at 1×10⁶ cells/well (2 mL/well) or 10 cm dishes at 9×10⁶ cells/dish (10 mL/dish), and cultured them for five days to reach 80-95% confluency. One day before stimulation, we incubated the cells overnight in DMEM containing 2.8 mM glucose, 2% BSA, 50 μM 2-mercaptoethanol, 10 mM nicotinamide, 5.5 μg/mL transferrin, 6.7 ng/mL sodium selenite, and 2 mM GlutaMAX. The next day, we washed the cells twice with HEPES-buffered Krebs-Ringer Buffer (KRB: 460 mM NaCl, 96 mM NaHCO₃, 20 mM KCl, 4 mM MgCl₂, 4 mM CaCl₂·2H₂O, 0.2% BSA, 10 mM HEPES, pH 7.4), and incubated them in KRB with 0.5 mM glucose for 60 minutes. After 1 hour, we removed the KRB solution and collected 100 µl aliquots for ELISAs. We added fresh KRB containing either 2 mM or 15 mM glucose to the cells for 1 hour and then collected 100 μL of supernatant to measure secreted insulin using the human insulin ELISA kit (Mercodia) and harvested RNA from the cells for downstream RNA analysis.

We calculated the insulin stimulation index as the ratio of the mean insulin content in the high glucose condition compared to the low glucose condition. We computed the 95% confidence interval of the stimulation index using Fieller’s method^27,28^ and *P*-values using the t-distribution, testing the null hypothesis that the stimulation index equals one.

### RNA isolation, poly(A) selection and quantitation

We harvested the cells from the 10 cm dishes by slowly mixing 1.5 mL of TRIzol^®^ reagent (TFS) into each sample on ice until the solution was homogenous. We aliquoted mixtures into 2 mL microfuge tubes, spun them at 3,000G for 5 min at 4°C, and stored them at -80°C. We isolated total RNA from the frozen TRIzol^®^ cell aliquots following the manufacturer’s instructions (TFS). We selected poly(A) RNA using NEBNext^®^ beads (NEB E3370S) and eluted with NEBNext^®^ Tris buffer (NEB E3370S) according to manufacturer’s instructions. We assessed the total RNA size distribution using the RNA 6000 Nano kit (Agilent) on the 2100 Bioanalyzer System (Agilent). We quantified each sample using Qubit® RNA BR Assay (TFS) per manufacturer’s instructions. We split the poly(A)-selected RNA samples into two aliquots, one for native RNA sequencing to retain the RNA modifications (native RNA), and one used to generate a matched IVT RNA sample without any RNA modifications (IVT RNA). For EndoC-BH1, we had sufficient RNA for one IVT sample in the high and low glucose conditions. We stored all samples at -80°C.

### cDNA amplification

From the IVT RNA aliquot, we performed first strand cDNA synthesis using the NEB template switching reverse transcriptase following the manufacturer’s instructions with the RT primer (5’-T_30_VN-3’) and template switching oligo (TSO; 5’-ACTC<u>TAATACGACTCACTATAG</u>GGAGAGGGCrGrGrG-3’; T7 promoter sequence is underlined) (NEB M0466). We PCR amplified the first strand cDNA products for 15 cycles at 98°C for 10 seconds, 65°C for 15 seconds, and 72°C for 3 minutes (NEB M0494). We purified the amplicons with AMPure XP Beads (Beckman Coulter™) according to the manufacturer’s protocol and quantified the cDNA using Qubit® DNA BR Assay (TFS). We concentrated each sample with a speed vacuum (Savant DNA120), saving 500 pg to confirm length distribution measurement using the High Sensitivity DNA kit (Agilent) on the 2100 Bioanalyzer (Agilent), and 1 μg for the IVT RNA synthesis reaction.

### *In vitro* transcribed RNA synthesis, purification, and quantitation

We used 1 µg of our cDNA template to synthesize IVT RNA using the HiScribe^®^ T7 High Yield RNA Synthesis Kit (NEB E2040S) according to manufacturer’s protocol, except that the samples were incubated for 2 hours at 42°C instead of 37°C. To reduce viscosity for purification, we diluted the IVT RNA reaction volume to 100 mL with 80 mL of RNAse-free 1X TE pH 8.0 (Quality Biological). We purified the IVT RNA using an equal volume of buffer-saturated phenol (pH 7.4):chloroform:isoamyl alcohol (25:24:1) followed by 2 pure chloroform washes. We performed an ethanol precipitation with 2.5X volumes of 100% ethanol and 0.1X volumes of 3M sodium acetate (pH 5.2), overnight storage at -20℃ and two washes with freshly made 70% ethanol. We resuspended the pellet in 50 mL of UltraPure™ DNAse/RNAse-Free Distilled Water (TFS). We quantified each RNA sample with a Qubit® RNA BR Assay, and assessed the size distribution with a RNA 6000 Nano Chip on a 2100 Bioanalyzer, as mentioned previously.

### Extension of IVT RNA 3’ ends with poly(A), purification, and quantitation

To ensure all IVT RNA molecules were adaptable for Direct RNA nanopore sequencing (dRNA-seq), we opted to *in vitro* poly(A) tail the IVT RNA using E. coli Poly(A) Polymerase (NEB M0276) according to manufacturer’s instructions. For each sample, we saved a 1.2 μL aliquot for a later RNA 6000 Nano Bioanalyzer analysis. Lastly, we purified the poly(A)-tailed IVT RNA with both MEGAclear™ Transcription Clean-Up Kit (TFS AM1908) and NEBNext^®^ beads following manufacturer’s protocol. After each subsequent clean-up procedure, we set aside 2.2 μL for Qubit® RNA BR Assay quantification and RNA 6000 Nano Chip Bioanalyzer analysis, as described previously, to assess pre- and post-poly(A) enrichment.

### Direct RNA nanopore sequencing and processing

We constructed sequencing libraries for each sample with 300 ng input poly(A) selected RNA using SQK-RNA004 sequencing kits following the manufacturer’s instructions (ONT). We loaded each library onto a FLO-PRO004RA flow cell and sequenced for 96 hours on a PromethION P24 running MinKNOW (v24.02.10). We basecalled the resulting pod5 files with dorado (v0.8.0) using the default read quality filters and the following models: rna004_130bps_sup@v5.1.0, rna004_130bps_sup@v5.1.0_inosine_m6A@v1, rna004_130bps_sup@v5.1.0_pseU@v1, and rna004_130bps_sup@v5.1.0_m5C@v1^29^. We aligned the reads from the unaligned bam files to gencode v46 reference transcriptome^30^ using dorado aligner (v0.8.0), filtered for forward primary alignments, and sorted with samtools (v1.17)^31^. We extracted the RNA modification calls from the bam files into bed files using modkit (v0.2.2)^32^. To quantify gene and transcript abundance, we created fastq files from the unaligned bam files, aligned to GRCh38 reference genome using minimap2 (v2.26) in splice-aware mode^33^, and filtered for primary alignments using samtools (v1.17) and isoquant^34^.

### Differential modification analysis

For each type of RNA modification, we tested for site-specific modifications associated with glucose exposure using binomial regression. For each cell line and modification site, we modeled the modification ratio by glucose condition using lme4 v1.1_35.5^35^ and the function glmer with options formula=”cbind(modified counts, unmodified counts) ∼ glucose condition”, family=”binomial”, control=lme4::glmerControl(optCtrl = list(iter.max = 1e5, eval.max = 1e5)), and the remaining parameters as default. We only considered modifications with a mean total count ≥20 and modification ratio ≥5% across the relevant cell line, as well as a modification ratio <5% in the IVT samples. To prune associations driven by outlier samples, we calculated Cook’s distances for each sample using the cooks.distance function from stats v4.3.3, derived *P*-values from the associated F-distribution, and removed any modification where the minimum Cook’s *P*-value <0.05. Next, we meta-analyzed modifications that were tested in both cell lines using the inverse-weighted fixed effects model as implemented by the rma function in metafor v4.6_0^36^ with default parameters. To focus on common effects across both cell lines, we removed any modification with evidence of heterogeneity across cell lines, defined as Cochran’s Q *P*-value <0.05. Finally, we performed multiple hypothesis correction using the Benjamini-Hochberg procedure^37^ and considered sites with a FDR<5% to be differentially modified. To account for standard errors in visualizations of effect sizes, we employed adaptive shrinkage using the function ash from ashr v2.2-63^38^ with the meta-analysis effect sizes and standard errors as input and the option mixcompdist="normal".

To test for enrichment, we used predicted T2D effector genes, annotated as moderate to causal, from the T2D knowledge portal^19,20^. For each modification and gene, if the gene contained a differentially modified site (FDR<5%), we labelled that gene as a success and performed a hypergeometric test (phyper function from stats v4.3.1 R package) across all genes considered in the dataset. We performed multiple hypothesis correction using the Benjamini-Hochberg procedure^37^.

### Differential gene and transcript expression analysis

For each cell line, we performed differential gene and transcript expression analysis to test for glucose-associated changes. We modeled the gene or transcript expression data from the isoquant^34^ quantification by glucose condition using DESeq2 (v1.36.0)^39^ with options test=”LRT”, fitType=”parametric”, reduced=”∼ 1”, sfType=”poscounts”, and the remaining parameters as defaults. Similar to the differential modification analysis, we only considered genes or transcripts with a mean total count ≥20 and a minimum Cook’s *P*-value <0.05. We meta-analyzed the genes or transcripts in both cell lines using the rma function in metafor v4.6_0^36^ with default parameters and removed those with evidence of heterogeneity across cell lines, indicated by Cochran’s Q *P*-value <0.05. Finally, we performed multiple hypothesis correction using the Benjamini-Hochberg procedure^37^ and considered features with a FDR<5% to be differentially expressed.

### Gene ontology enrichment and clustering analysis

We calculated the enrichment of the differentially modified or expressed genes (FDR<5%). We used the biological process gene ontology (GO) terms and the clusterProfiler v4.8.3^40^ package compareCluster function with enrichGO as the enrichment function. To perform multiple hypothesis correction, we used the Benjamini-Hochberg procedure^37^.

We visualized the enrichment results (FDR<5%) using the emapplot function from the enrichplot v1.20.3^41^ package to create networks where vertices depict GO terms and edges connect similar terms. Briefly, we used the godata function from GOSemSim v2.26.1^42^ to model the semantic similarity between GO terms and the pairwise_termsim function from enrichplot with the method parameter “Rel” to calculate the similarity matrix. We generated plots of the similarity matrix using the the emapplot function with the following options: node_label=“group”, edge.params=list(min=0.75), pie.params=list(pie=“Count”), and cluster.params=list(cluster=T). The edges connect GO terms (vertices) with a similarity ≥0.75, the pie charts represent the relative proportion of genes within a GO term, and terms are grouped into clusters based on their shared, semantic similarity (clusters calculated with kmeans function from stats v4.3.2 where k=sqrt[number_vertices] as implemented in emapplot). We labeled each group of terms manually based on the GO terms within each group.

## Code availability

The differential modification code is available as a snakemake pipeline at https://github.com/CollinsLabBioComp/snakemake-binomial_regression.

## Funding

This work was supported by the US National Institutes of Health (NIH) grants 1ZIAHG000024-30 (to FSC and LGB) and 1K99DK139175-01 (to DLT), the NIH Intramural Sequencing Center grant 1ZIBHG000196 (to DLT, FSC, and LGB), the Italian Association for Cancer Research (AIRC) project IG 2019 ID 22851 (to FN), the European Molecular Biology Laboratory (to EB), the EMBL ETPOD fellowship (to LM), the Gates Cambridge Trust (to HJT), and the NIH Oxford-Cambridge scholars’ programme (to HJT).

## Author contributions

LM, EB, FSC, and DLT designed the research. AL, AJS, LLB, SYB, LGB, MRE, FSC, and DLT performed research. LM, HJT, MZ, BNL, TF, NN, FN, EB, and DLT analysed data. LM, EB, FSC, and DLT supervised the study. LM, HJT, LGB, EB, FSC, and DLT wrote the paper. All authors reviewed and approved the paper.

## Conflicts of interest

LM has received reimbursement of travel or accommodation expenses to speak at Oxford Nanopore Technologies (ONT) conferences. EB is a paid consultant and shareholder of ONT.

## Supplemental Tables

**S Table 1.** Sequencing Summary Statistics. Sample-level summary statistics. Genes and transcripts detected with *≥* 3 reads. Samples denoted by cell line, native or IVT, glucose condition, and replicate number.

| Samples | Number of Reads (M) | Percent Aligned Reads | Transcripts Detected | Genes Detected | N50 (kb) | Mean Read Length | Median Read Length |
| --- | --- | --- | --- | --- | --- | --- | --- |
| BH1_native_H_r1 | 19.65 | 99.96 | 40254 | 17372 | 1.79 | 1267.05 | 921 |
| BH1_native_H_r2 | 20.57 | 99.94 | 41016 | 17578 | 1.79 | 1274.18 | 934 |
| BH1_native_H_r3 | 18.61 | 99.94 | 40258 | 17447 | 1.77 | 1253.05 | 912 |
| BH1_native_L_r1 | 21.77 | 99.94 | 42129 | 17747 | 1.80 | 1284.09 | 940 |
| BH1_native_L_r2 | 21.77 | 99.94 | 41050 | 17517 | 1.80 | 1284.09 | 940 |
| BH1_native_L_r3 | 21.77 | 99.94 | 41964 | 17615 | 1.80 | 1284.09 | 940 |
| BH3_native_H_r1 | 17.18 | 99.94 | 35605 | 16013 | 1.59 | 1104.12 | 780 |
| BH3_native_H_r2 | 15.62 | 99.97 | 33826 | 15537 | 1.74 | 1212.03 | 860 |
| BH3_native_H_r3 | 15.58 | 99.97 | 34055 | 15626 | 1.74 | 1220.17 | 863 |
| BH3_native_H_r4 | 18.38 | 99.96 | 34966 | 15827 | 1.71 | 1197.75 | 856 |
| BH3_native_L_r1 | 16.10 | 99.92 | 34925 | 15652 | 1.67 | 1156.41 | 821 |
| BH3_native_L_r2 | 15.20 | 99.97 | 32247 | 15116 | 1.70 | 1191.79 | 851 |
| BH3_native_L_r3 | 9.67 | 99.96 | 29300 | 14430 | 1.73 | 1196.87 | 844 |
| BH3_native_L_r4 | 19.62 | 99.96 | 31957 | 15069 | 1.64 | 1137.10 | 800 |
| BH1_IVT_H_r1 | 2.06 | 99.76 | 18936 | 9079 | 0.475 | 436.68 | 366 |
| BH1_IVT_L_r1 | 2.10 | 99.72 | 19415 | 9226 | 0.467 | 427.84 | 358 |
| BH3_IVT_H_p1 | 1.46 | 99.84 | 17748 | 8223 | 0.514 | 446.79 | 376 |
| BH3_IVT_H_p2 | 1.96 | 99.83 | 19711 | 8764 | 0.519 | 443.32 | 360 |
| BH3_IVT_H_r1 | 4.19 | 99.66 | 24744 | 10201 | 0.438 | 384.76 | 311 |
| BH3_IVT_H_r2 | 3.12 | 99.67 | 21752 | 9341 | 0.465 | 405.82 | 326 |
| BH3_IVT_H_r3 | 1.04 | 99.36 | 14229 | 6863 | 0.413 | 356.70 | 289 |
| BH3_IVT_L_r1 | 3.29 | 99.77 | 20306 | 8818 | 0.534 | 468.09 | 386 |
| BH3_IVT_L_r2 | 3.46 | 99.82 | 21347 | 9155 | 0.493 | 439.39 | 367 |
| BH3_IVT_L_r3 | 3.35 | 99.78 | 20686 | 8894 | 0.477 | 422.08 | 347 |

**S Table 2.** RNA modification effector DGE and DTE results. Gene or transcript expression changes for known RNA modification writer, reader, and eraser genes. We report the minimum *P*-value and *q*-value from the differential expression results for the gene or any of the corresponding transcripts.

| Gene Symbol | Effector Type | Modification | Minimum Gene *P*-value | Minimum Gene *q*-value |
| --- | --- | --- | --- | --- |
| ADAR | writer | inosine | 0.091573 | 0.849601 |
| NSUN2 | writer | m5C | 0.230276 | 0.808251 |
| NSUN6 | writer | m5C | 0.839333 | 0.996113 |
| ALYREF | reader | m5C | 0.439041 | 0.907533 |
| RAD52 | reader | m5C | 0.254160 | 0.825605 |
| SRSF2 | reader | m5C | 0.001604 | 0.141133 |
| YBX1 | reader | m5C | 0.019135 | 0.525202 |
| ALKBH1 | eraser | m5C | 0.175732 | 0.937246 |
| TET1 | eraser | m5C | 0.049190 | 0.553876 |
| TET2 | eraser | m5C | 0.737255 | 0.970146 |
| TET3 | eraser | m5C | 0.626204 | 0.955300 |
| METTL14 | writer | m6A | 0.278299 | 0.858438 |
| METTL3 | writer | m6A | 0.029178 | 0.529414 |
| RBM15 | writer | m6A | 0.267021 | 0.828117 |
| RBM15B | writer | m6A | 0.011761 | 0.285843 |
| WTAP | writer | m6A | 0.186124 | 0.941964 |
| HNRNPA2B1 | reader | m6A | 0.014807 | 0.487587 |
| HNRNPC | reader | m6A | 0.038105 | 0.681313 |
| SRSF10 | reader | m6A | 0.056839 | 0.751188 |
| SRSF3 | reader | m6A | 0.283413 | 0.836723 |
| YTHDC1 | reader | m6A | 0.032434 | 0.456420 |
| YTHDC2 | reader | m6A | 0.694808 | 0.963389 |
| YTHDF1 | reader | m6A | 0.970352 | 0.997533 |
| YTHDF3 | reader | m6A | 0.123122 | 0.892943 |
| ALKBH5 | eraser | m6A | 0.132907 | 0.712594 |
| FTO | eraser | m6A | 0.661733 | 0.959751 |
| YTHDF2 | reader | m6A & m5C | 0.176247 | 0.757396 |
| PUS1 | writer | pseudouridine | 0.060925 | 0.576212 |
| PUS10 | writer | pseudouridine | 0.595974 | 0.986601 |
| PUS3 | writer | pseudouridine | 0.698064 | 0.992503 |
| PUS7 | writer | pseudouridine | 0.235122 | 0.970057 |
| PUS7L | writer | pseudouridine | 0.312016 | 0.847353 |
| PUSL1 | writer | pseudouridine | 0.047473 | 0.529734 |
| RPUSD1 | writer | pseudouridine | 0.282757 | 0.988666 |
| RPUSD2 | writer | pseudouridine | 0.734146 | 0.969895 |
| RPUSD3 | writer | pseudouridine | 0.075340 | 0.619455 |
| RPUSD4 | writer | pseudouridine | 0.231712 | 0.808808 |
| TRUB1 | writer | pseudouridine | 0.543655 | 0.937439 |
| PFN1 | reader | pseudouridine | 0.033006 | 0.461556 |

## Supplemental Figures

**S Fig. 1.**
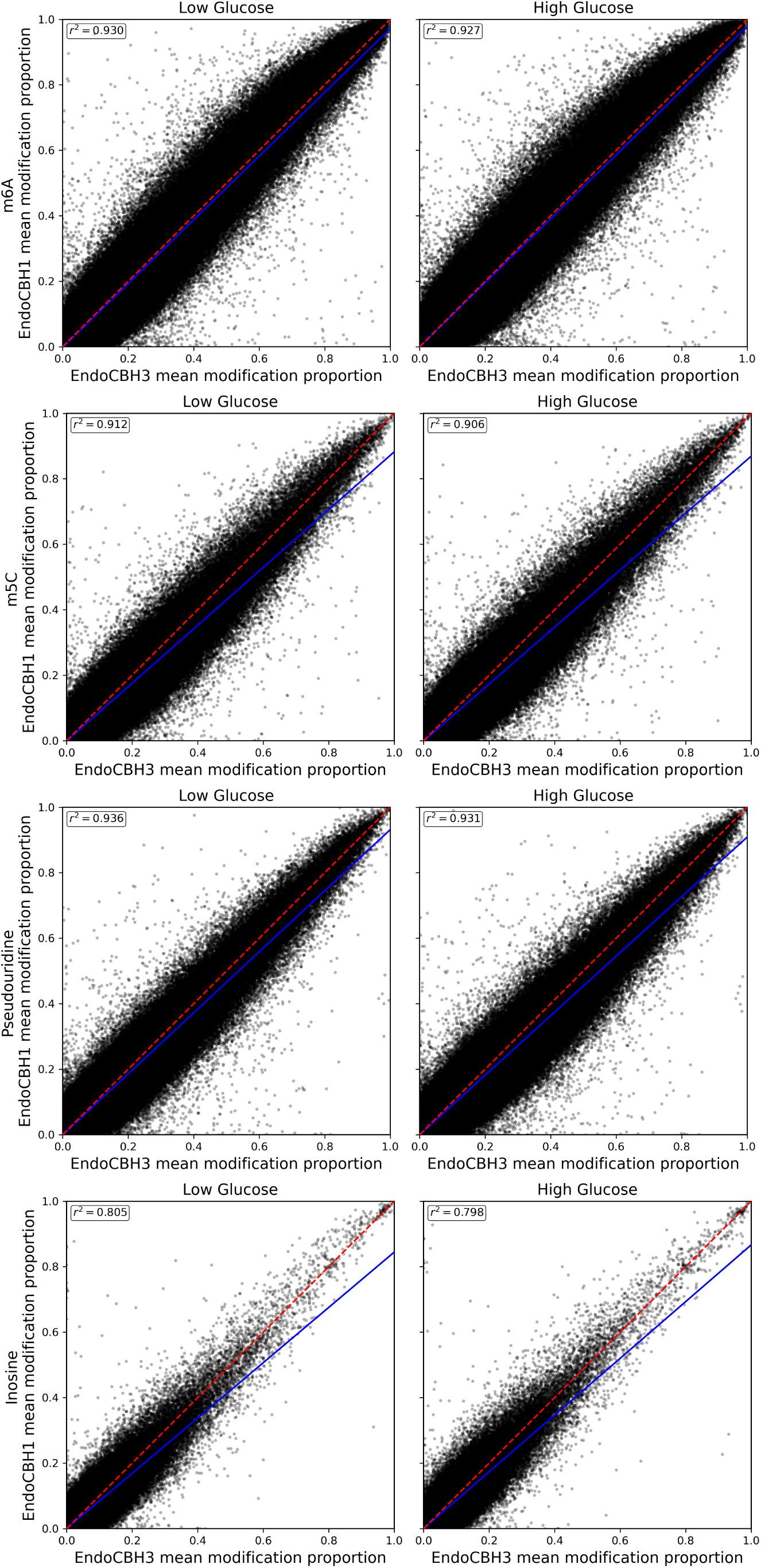
Correlation of EndoC-BH1 and EndoC-BH3 modification proportion. Correlation of average RNA modification proportion between EndoC-BH1 (y-axis) and EndoC-BH3 (x-axis) for low and high glucose conditions (facets x-axis) and for the RNA modifications considered (facets y-axis). Dashed red line corresponds to the identity line; the blue line corresponds to the least squares regression line.

**S Fig. 2.**
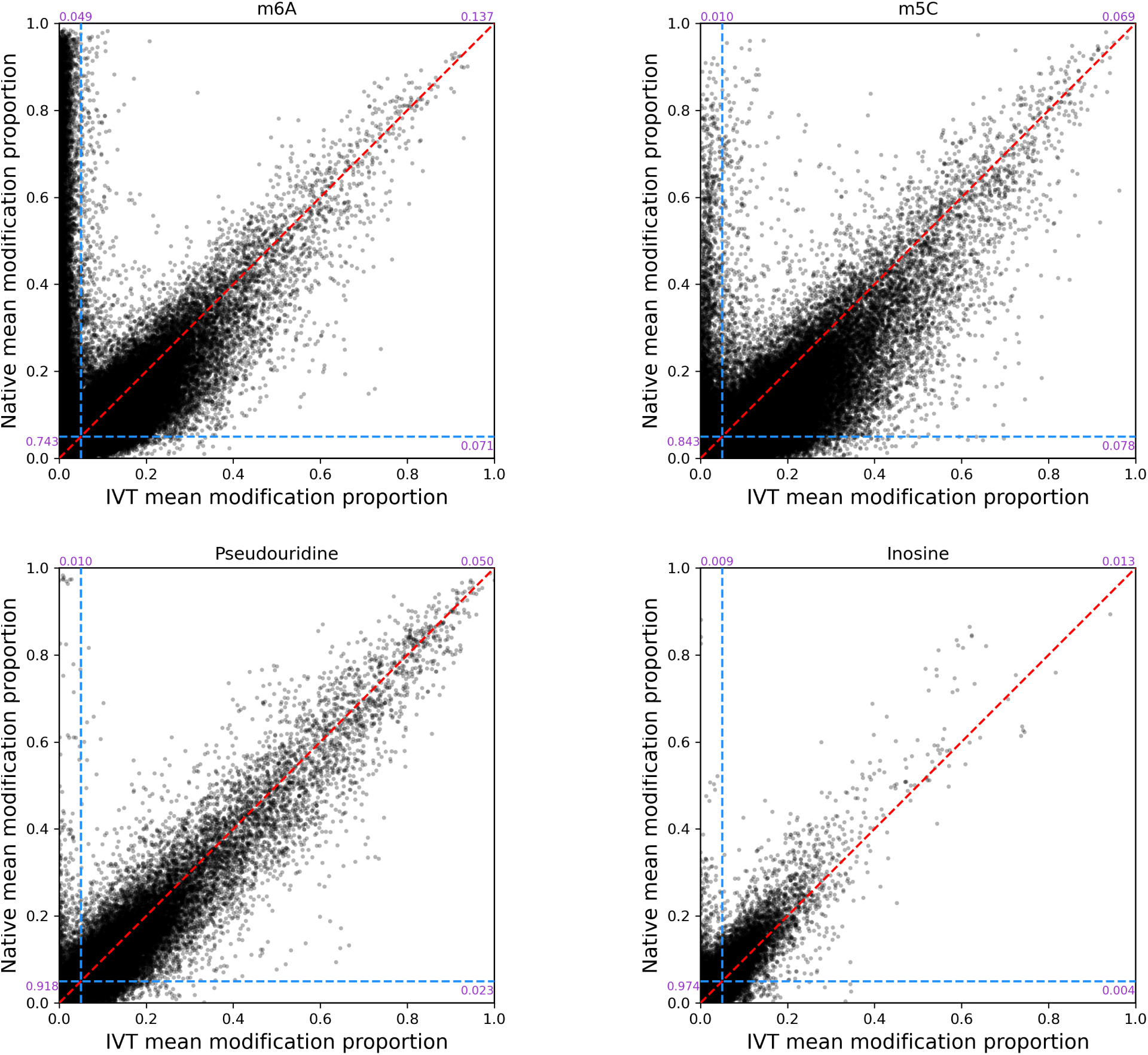
Correlation of Native and IVT modification proportion. Correlation of average RNA modification proportion between native (y-axis) and IVT (x-axis) for the RNA modifications considered (facets). Red line corresponds to the identity line; dashed blue lines depict the filtering thresholds. The purple values report the proportion of points within each quadrant defined by the blue, filtering threshold lines.

**S Fig. 3.**
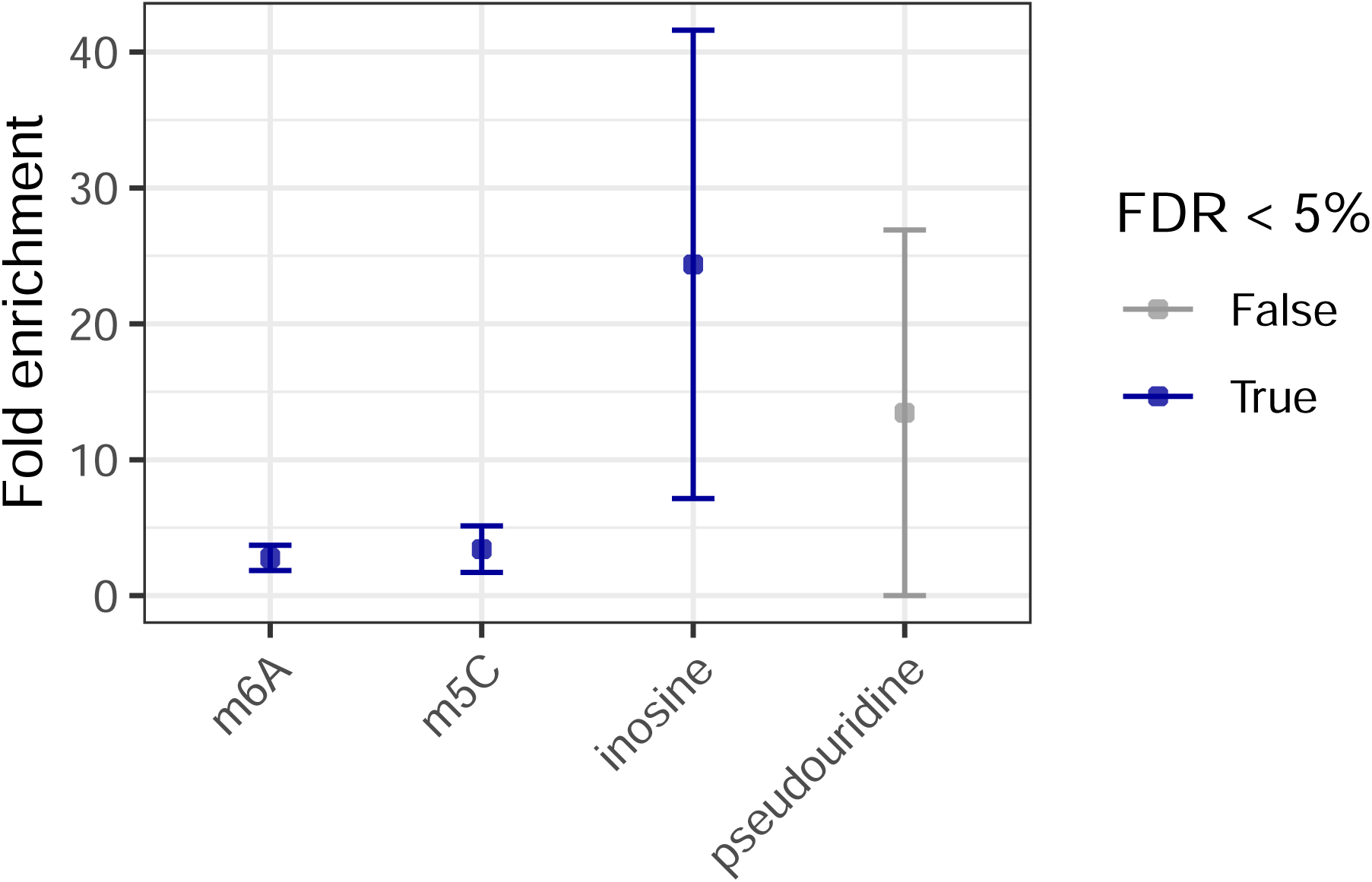
Enrichment of known T2D genes. The fold enrichment for T2D genes in the DMSs (y-axis) for the RNA modifications considered (x-axis). Color indicates FDR*<* 5%.

**S Fig. 4.**
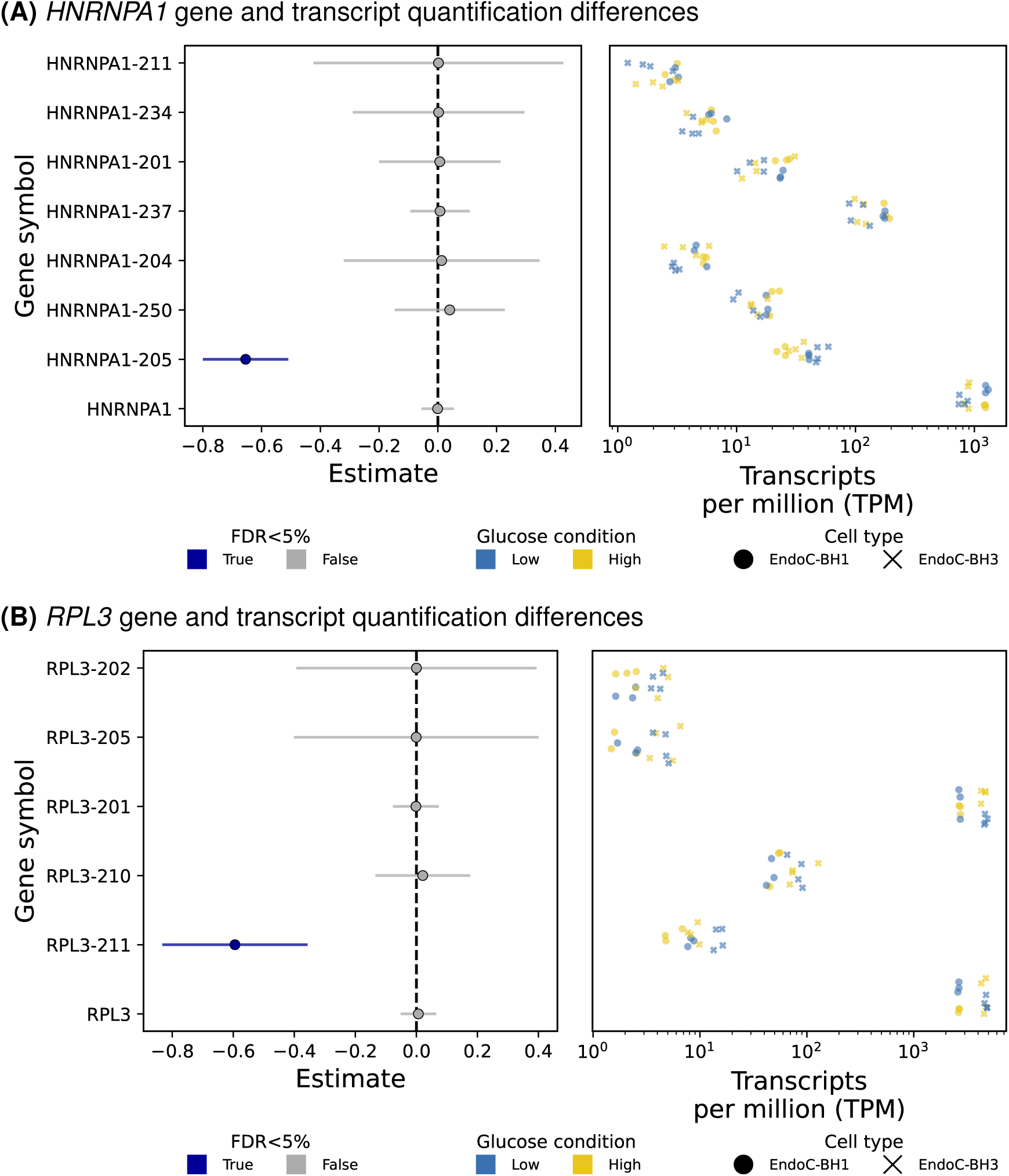
Gene and transcript effects. Forest plot (left) and corresponding sample transcripts per million (TPM) (right) for (A) *HNRNPA1* and (B) *RPL3*. The forest plot shows the effect size estimate (x-axis) for each transcript (y-axis). FDR*<* 5% shown by blue points and the 95% confidence interval represented by the error lines. The right plot shows the TPM values (x-axis) for the corresponding gene or transcript (y-axis) across samples (points), colored by glucose condition. Point shape distinguishes cell line: EndoC-BH1 (circle) or EndoC-BH3 (X).

**S Fig. 5.**
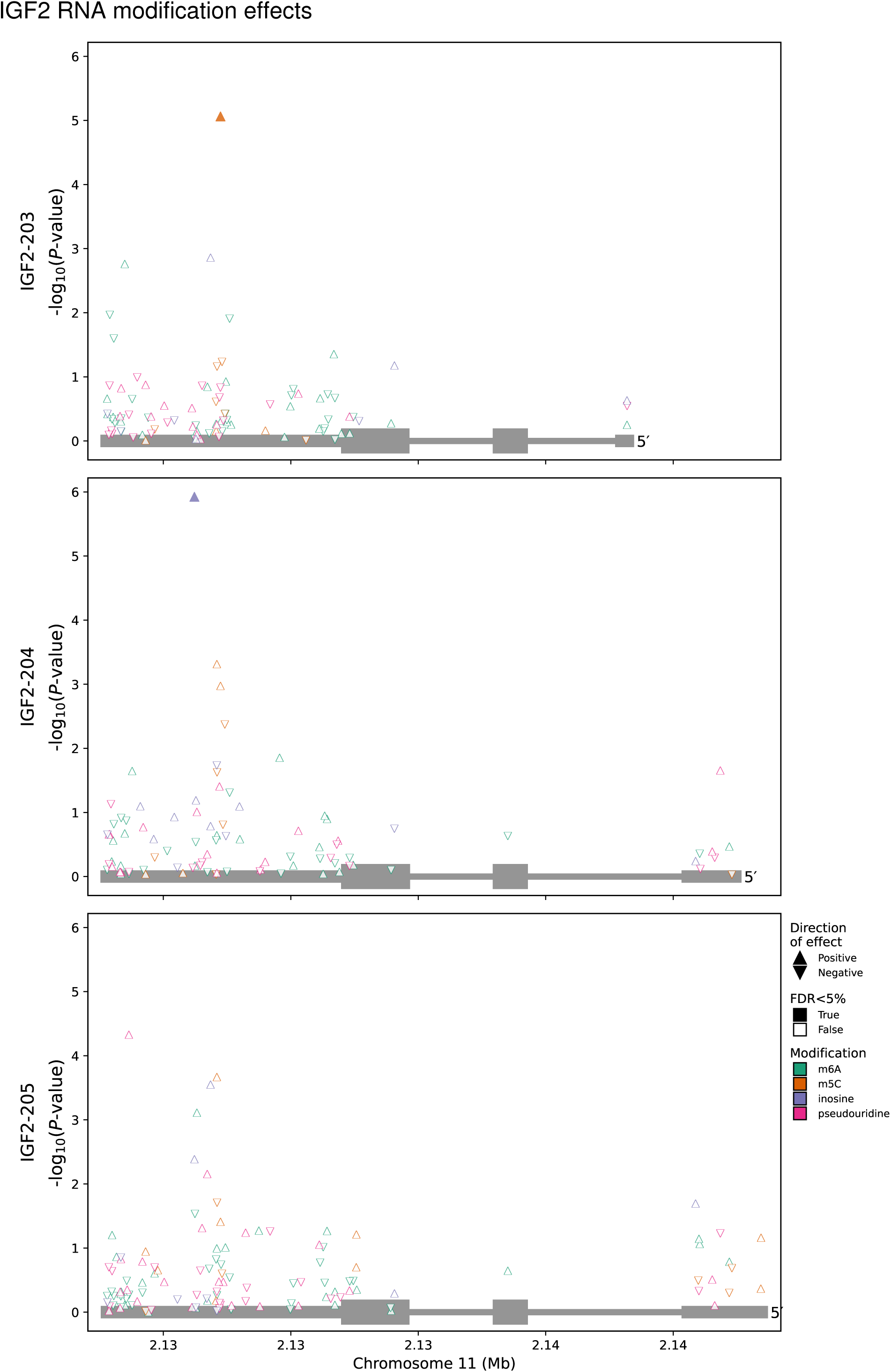
*IGF2* RNA modification results at transcript resolution. RNA modification sites considered (points) along with their genomic coordinates (x-axis) in relationship to the detected transcripts (y-axis facets) of *IGF2* (grey boxes) and the differential modification *−log*_10_(*P*-value) (y-axis). Direction of effect indicated by triangle orientation, where an upward triangle indicates a positive effect (i.e., increased proportion of modified counts in the high glucose condition). Colors depict the RNA modification. Fill status indicates FDR*<* 5%. GO terms enriched in differential expression results

**S Fig. 6.**
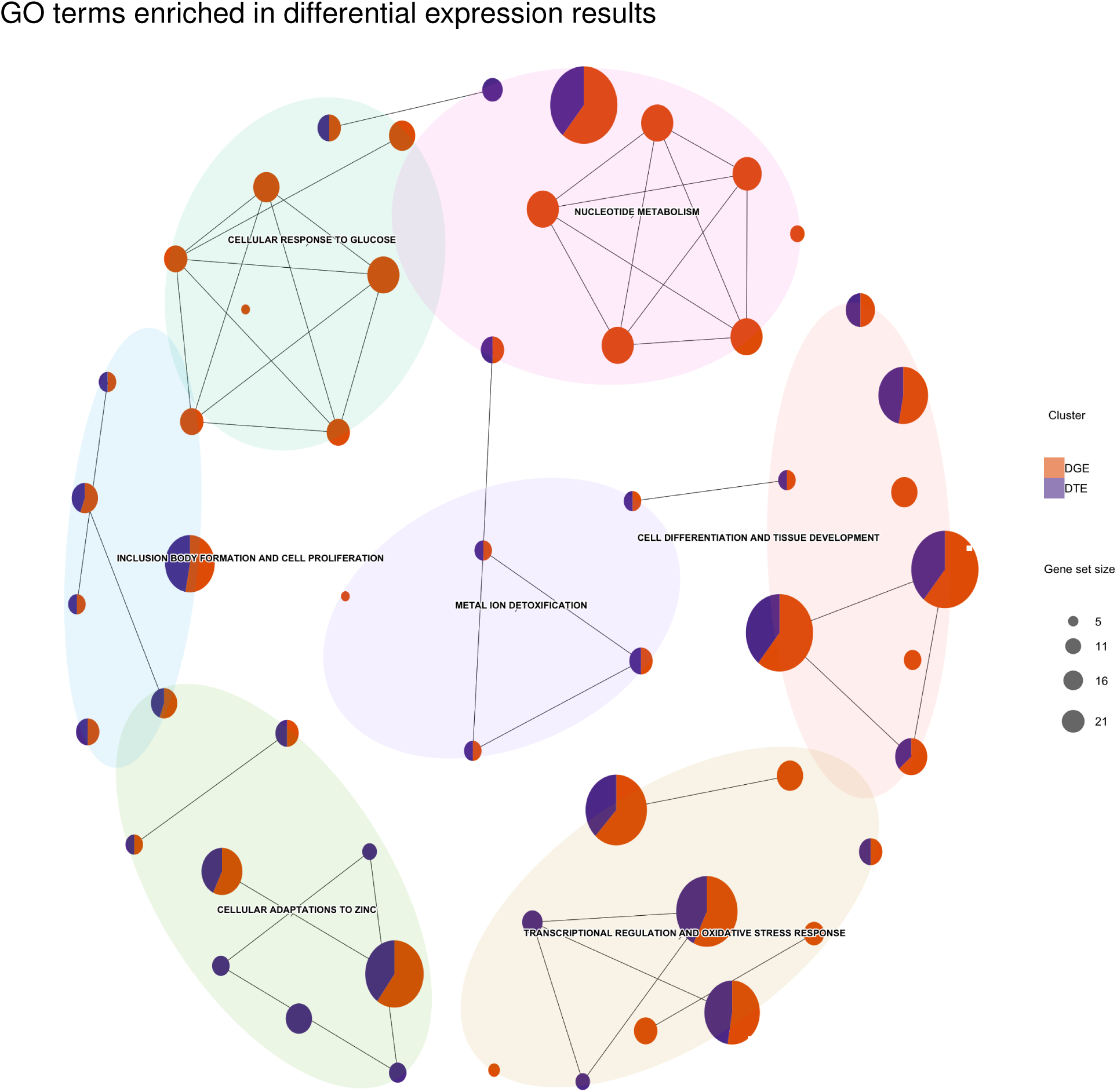
GO enrichments for differential gene and transcript expression results. Network of GO terms enriched (FDR*<* 5%) for differential gene and transcript expression. Each node is a GO term with the proportion of genes from DGE (orange) or DTE (purple) represented by the pie charts. The node size is scaled by the number of genes overlapping each GO term. Edges connect semantically similar nodes, which are also clustered based on similarity.

